# Eccentric cycling improves motor learning more than concentric cycling

**DOI:** 10.1101/2025.05.08.652761

**Authors:** Layale Youssef, Benjamin Pageaux, Jason L. Neva

**Author notes:** ***Corresponding Author:*** Layale Youssef, 4545 Queen Mary Road, Montreal, QC, Canada. Both authors equally contributed to this work (Benjamin Pageaux, Jason L. Neva).

## Abstract

An acute bout of aerobic exercise (AAE) performed before practicing a motor task can enhance skill acquisition and motor learning. To date, research on the effects of AAE on motor learning has focused exclusively on concentric cycling, leaving the impact of eccentric cycling unexplored. Unlike concentric cycling, eccentric cycling involves muscle lengthening while resisting the reverse movement of the pedals and is characterized by greater force production with lower cardiovascular and metabolic cost. Regarding neuroplasticity changes, eccentric contractions induced a prolonged decrease in intracortical inhibition compared to concentric contractions. Eccentric cycling AAE also increases activation in cognitive-related regions. Given the involvement of these regions and motor cortex excitability in motor learning, we hypothesized that eccentric cycling AAE would enhance motor learning to a greater extent than concentric AAE. A total of 60 young healthy individuals were allocated to one of three groups that performed 20 min of: *i)* eccentric cycling; *ii)* concentric cycling; or *iii)* seated rest. Both cycling AAE conditions were performed at a power equivalent to 70% peak heart rate (i.e., moderate intensity). A continuous tracking task was used to assess motor skill acquisition (immediately after the intervention) and motor learning (48 h retention test). For both acquisition and retention, the eccentric group outperformed both the concentric and rest groups, while the concentric group also showed a better performance compared to the rest group at retention. Thus, we demonstrated that eccentric cycling AAE enhances motor learning to a greater extent than concentric cycling AAE, while also confirming previous work that showed enhanced motor learning following concentric cycling AAE compared to rest. Our findings suggest that eccentric cycling AAE may have important implications for exercise protocols prescribed in sports-related and clinical contexts.

**Highlights:**

- Eccentric cycling enhanced motor learning to a greater extent than concentric cycling
- Eccentric and concentric cycling enhanced motor learning more than rest
- Enhanced skill acquisition occurred after eccentric cycling, along with lower heart rate response and perceived effort.
- Eccentric cycling may have important implications in sports-related and clinical contexts

## Introduction

A single session of physical exercise can enhance cognitive performance (i.e., processing, flexibility, decision-making) and learning motor skills ^1-5^. More specifically, an acute bout of aerobic cycling exercise (AAE) performed close in time to practicing a motor task can enhance skill acquisition ^6-8^ and promote motor learning ^4^, demonstrated by improved performance at retention tests 5 h ^9^, 24 h ^6,8,10-12^ and 7 days ^10,11,13,14^ after practice. The impact of AAE on motor learning has been demonstrated by manipulating different exercise parameters, such as intensity (e.g., low, moderate, high), type (e.g., continuous, interval) and duration (e.g., 5-35 min) ^4,6,8,9,15^. Critically, all work to date demonstrating the impact of AAE on motor learning have used the traditional form of muscle contraction during exercise, which is referred to as concentric cycling ^8,16-19^. This traditional form of lower-limb cycling involves cycling forward by contracting the muscles of the lower limbs, causing the muscle fibers to shorten to generate movement ^20,21^. To date, no study has investigated the potential of cycling AAE to increase motor learning using a different modality of muscle contraction (i.e., the eccentric contraction mode).

Eccentric cycling is a mode of lower-limb cycling that is less conventional as it involves contracting the quadriceps muscles while resisting force produced by a motor generating a backwards cycling motion, where muscle fibers are lengthened to produce movement ^22^. Compared to concentric cycling, eccentric cycling presents the advantage of producing a higher level of power output while requiring a lower cardiovascular, respiratory and metabolic cost ^23-26^. Eccentric cycling interventions have generated promising applicability in various populations, such as patients with cardiorespiratory limitations ^27^ and in the aging population ^28^. Eccentric exercise, specifically downhill treadmill exercise, has been shown to elicit a similar increase in corticospinal excitability in the non-exercised upper limbs as concentric exercise, which involved uphill treadmill walking ^29^. Additionally, prolonged decrease of intracortical inhibition of the primary motor cortex following eccentric contractions have been observed ^30^. Other research has shown that eccentric cycling AAE enhances activity in the bilateral prefrontal cortices and the right parietal cortex to a greater extent than concentric cycling AAE ^31^. Considering the importance of prefrontal cortical activity ^32^ and motor cortex excitability changes during the process of motor learning ^32-36^, it is possible that eccentric cycling AAE may enhance motor learning to a greater extent than traditional concentric cycling AAE ^6,16,37,38^.

Thus, the objective of this study was to examine the impact of eccentric cycling AAE on motor learning. We hypothesized that eccentric cycling AAE performed prior to motor task practice will enhance motor sequence learning to a greater extent than concentric cycling AAE and rest. Additionally, based on previous literature ^8,38-40^, we expected that concentric cycling AAE would also improve skill acquisition and motor learning compared to rest.

## Methods

### Participants

A total of 60 young healthy individuals (30 men and 30 women) between the ages of 18 and 38 years old (26.2 ± 3.7) were recruited to participate in this study (Table 1). A sensitivity analysis performed in G*Power with an alpha risk of 0.05 and a sample size of 60 indicated that we had a 90% chance of observing a medium effect size of f(U) = 0.234 (∼ *η*^*2*^_*p*_ = <u>0.052</u>) or higher ^41^. Participants had no known neurological disorders and the Physical Activity Readiness Questionnaire (PAR-Q) indicated they were in sufficiently good health to complete the exercise protocols. All participants filled in the International Physical Activity Questionnaire (IPAQ) to estimate their physical activity level. The Aging and Neuroimaging Research Ethics Board of the Research Center of the Institut Universitaire de Gériatrie de Montréal (CRIUGM) approved all experimental procedures, and all participants provided written informed consent before participating in the study.

**Table 1:**
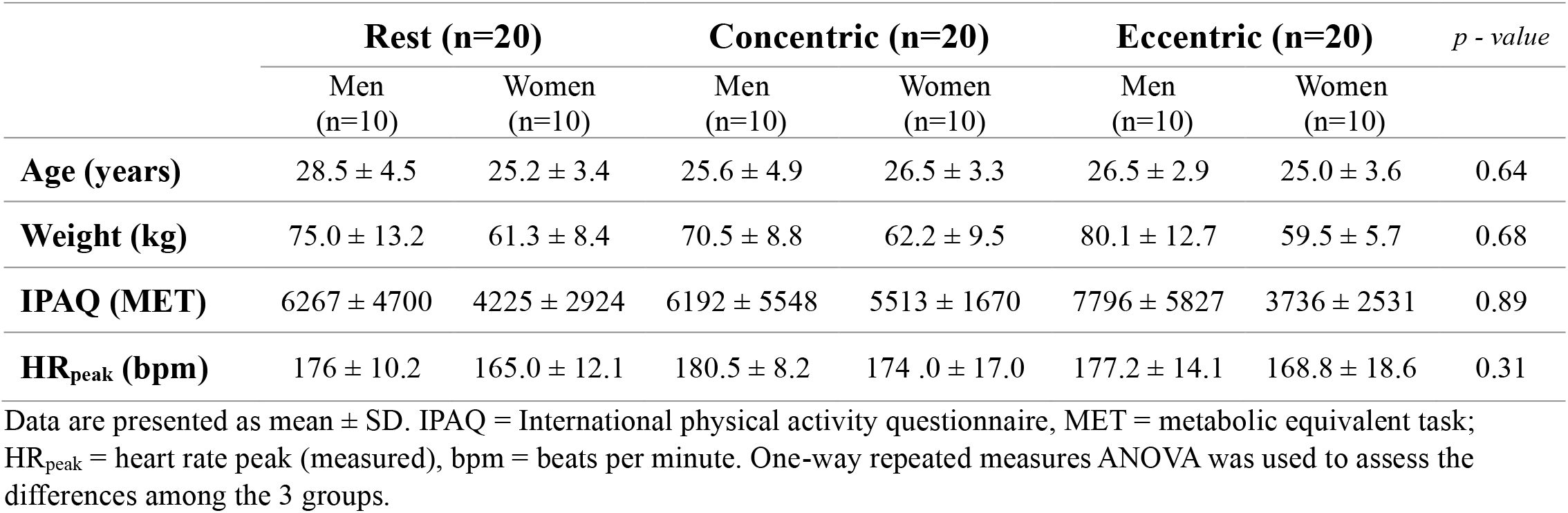
Participants’ characteristics.

### Experimental design

Participants were divided into three groups that corresponded to the following experimental interventions: 1) eccentric cycling exercise, 2) concentric cycling exercise and 3) seated rest. Each participant completed three visits over a maximum period of two weeks. Visit 1 consisted of an incremental exercise test and familiarisation with eccentric cycling that was completed by all participants regardless of experimental group allocation. Following this visit, participants were assigned to one of the three experimental groups. Visit 2 consisted of participating in 25 minutes of the exercise (eccentric or concentric) or rest intervention followed by motor task practice. Visit 3 consisted of a 48-h retention test of the motor task. An overview of the experimental design is presented in Figure 1.

**Figure 1:**
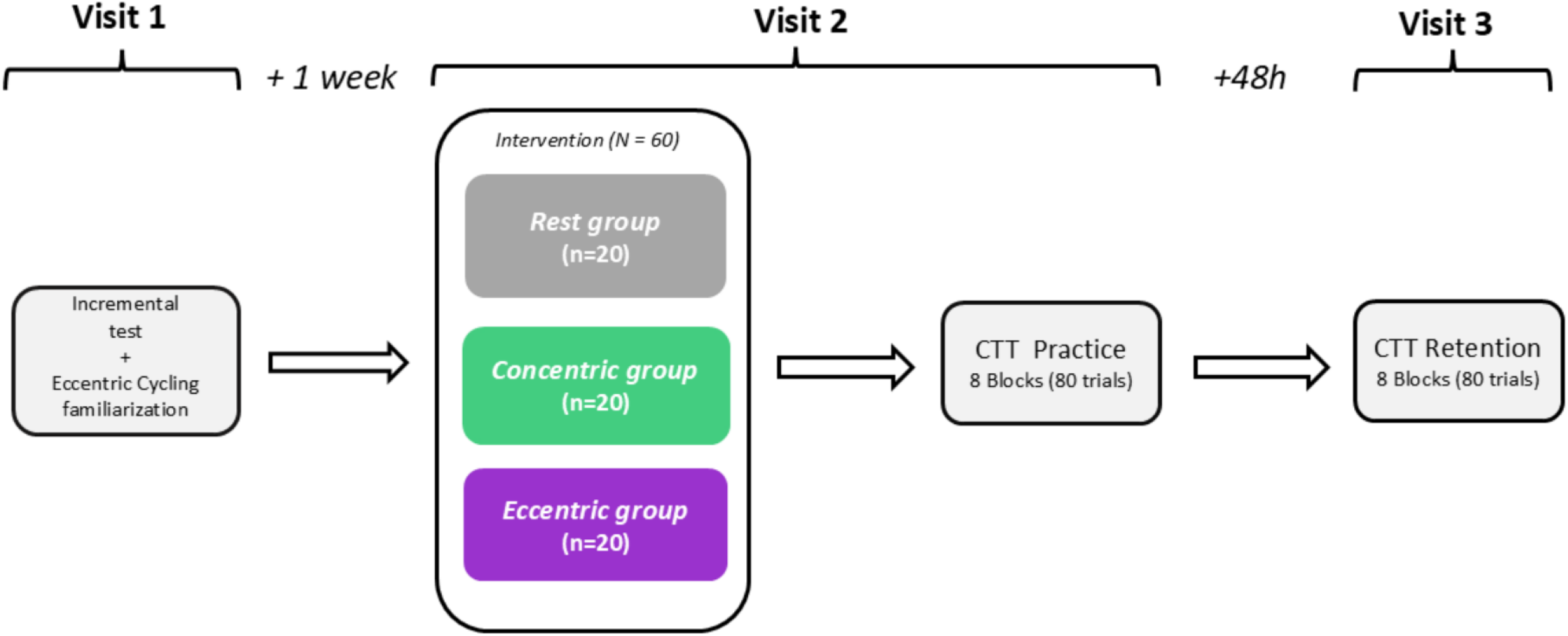
Study protocol overview. Visit 1 included an incremental test and a familiarisation with eccentric cycling for all participants. Visit 2 took place one week later, where participants were assigned to one of the three groups: Rest group, Concentric group, or Eccentric group, undergoing 20 minutes of rest, concentric cycling, or eccentric cycling, respectively. Immediately after the intervention, all participants completed 8 blocks of the CTT. Visit 3 took place 48 hours post-intervention and consisted of the CTT retention test (8 blocks). CTT (continuous tracking task) refers to the motor task.

### The incremental exercise test

An incremental exercise test was conducted on a concentric recumbent cycle ergometer (CY00500, Cyclus 2, Germany). The increments and the starting workload were determined based on each participant’s body weight ^42,43^. The increments increased every 2 min until exhaustion. Participants sat comfortably on a cycle ergometer and were asked to maintain a cycling cadence of 60 rotations per minute (rpm) throughout the test. During the test, heart rate was continuously monitored using a heart rate sensor (Polar H10, Polar Electro 2024, Finland) and heart rate peak (HR_peak_) was obtained. Perceived effort and muscle pain using the CR100 scale ^44,45^ as well as affect using the Feeling scale ^46^ were reported at the end of each increment (every 2 min).

### Acute Aerobic Exercise (AAE) and rest

The AAE intervention lasted 25 minutes, consisting of a 5-minute warm-up followed by 20 minutes of continuous moderate-intensity cycling. The warm-up was performed at 50% HR_peak_, and the exercise at 70% HR_peak_^43,47^. For both concentric and eccentric cycling, the power output was set to correspond to the power output eliciting 70% of the participant’s HR_peak_ during the incremental test previously described. During both eccentric (Cyclus 2, Germany; model CY00360) and concentric (Cyclus 2, Germany; model CY00500) cycling, the ergometers were set in isopower mode, and the participants were asked to maintain a cadence of 60 rpm. Concentric and eccentric ergometers were equipped with identical frames, i.e., the motor for concentric cycling (CY00500) was mounted on an identical bike frame of the eccentric ergometer. During both cycling exercises, heart rate was monitored and recorded every 5 minutes. Additionally, participants reported their perceived effort and muscle pain using the CR100 scale ^44,45^, as well as their affect using the Feeling scale ^46^.

The rest intervention lasted 25 minutes, where participants were comfortably seated on a chair watching an emotionally neutral documentary ^48^. Participants were specifically instructed to keep their upper limbs inactive. Heart rate, perceived effort, muscle pain and affect were reported every 5 minutes, similarly to the AAE interventions.

### The motor task

The motor task used in this study was the continuous tracking task (CTT), where participants manipulated a custom joystick using their dominant thumb. During the CTT, participants manipulated the movement of a cursor up and down to follow the vertical trajectory of a target using a custom joystick. At the same time, the target and cursor were always horizontally aligned and moved horizontally at a constant velocity from the right to the left on the computer screen. Participants were instructed to control the cursor to track the moving target as quickly and accurately as possible throughout the task. Joystick position sampling and stimulus presentation occurred at a frequency of 50 Hz, collected using custom software created with LabView (v. 9.0; National Instruments, Austin, TX). Each trial lasted 30 s and was preceded by a 2-s period of normalization (where the cursor aligned with the target vertically). The first and last 10 seconds of each trial consisted of random sequences, while the middle 10 seconds consisted of a repeated sequence that was consistent across all blocks during both the practice and retention phases. Improved performance at the retention test with random sequence practice reflects non-specific learning, while improved performance with repeated sequence practice reflects specific-sequence learning ^16,49-51^. Prior to the intervention (eccentric AAE, concentric AAE or rest), participants practiced 3 CTT trials to assess pre-intervention performance. Practice of the CTT was performed by all participants on Visit 2 immediately after the intervention (acquisition: 8 blocks of 10 trials, 80 trials total, taking ∼ 40 min) as well as on Visit 3 for a retention test which took place 48 h later (retention: 8 blocks of 10 trials, 80 trials total, taking ∼40 min).

The data from CTT practice were processed using a custom script in MATLAB (Mathworks, R2016a). For each sequence, root mean squared error (RMSE) was calculated. RMSE was calculated separately for the repeated sequence and the random sequences (average of the first 10 s and last 10 s of each trial). Higher RMSE values indicate less accurate target tracking (i.e., worse performance) and lower RMSE values indicated more accurate target tracking (i.e., better performance) during CTT practice. Outliers corresponding to RMSE values higher than 2 standard deviations for each block of practice for each participant were identified as outliers and were removed (corresponding to 0.8% of all trials).

## Statistical analyses

Two-way mixed model ANOVAs were conducted on physiological and psychological data measured during exercise or rest, specifically % HR_peak_, perceived effort, muscle pain, and affect, with between-subjects factor GROUP (eccentric, concentric, rest) and within-subjects factor TIME (5, 10, 15, 20 min). Since perceived effort and muscle pain were rated as zero in the rest group, this group was excluded from the analysis of these two variables, resulting in a direct comparison between the concentric and eccentric cycling exercise groups. For % HR_peak_ and affect, all three groups (rest, concentric, eccentric) were included in the analysis.

Three-way mixed model ANOVAs were conducted for the variable RMSE, with between-subjects factor GROUP (eccentric, concentric, rest) and within-subjects factors SEQUENCE (random, repeated) and BLOCK (all 8 blocks) on the CTT data. These analyses were performed separately for acquisition (motor practice immediately after the intervention) and retention (motor practice 48 hours after the intervention). For acquisition, we conducted an additional mixed model ANOVA using the calculated change score between block 8 and block 1 (B8-B1) to assess the magnitude of motor performance improvement, with between-subjects factor GROUP (eccentric, concentric, rest) and within-subjects factor SEQUENCE (random, repeated). Post-hoc analyses were performed using Bonferroni correction where appropriate. Results were considered statistically significant with a *p*-value of < .05. Effect sizes are reported as partial eta squared (η^2^_p_) for ANOVAs (0.01, 0.06, and 0.14 corresponding to small, moderate, and large effects, respectively), and as Cohen’s *d* for pairwise comparisons (0.2, 0.5 and 0.8 corresponding to small, moderate, and large effects, respectively)^52^. All statistical analyses were conducted using R software (4.2.1) (foundation for statistical computing, Vienna, Austria) using the packages *afex and stats*.

## Results

### Physiological and perceptual responses during the interventions

Heart rate, perceived effort, muscle pain and affect were reported every 5 min during the intervention (Figure 2 and Figure S1 in Supplementary material).

**Figure 2:**
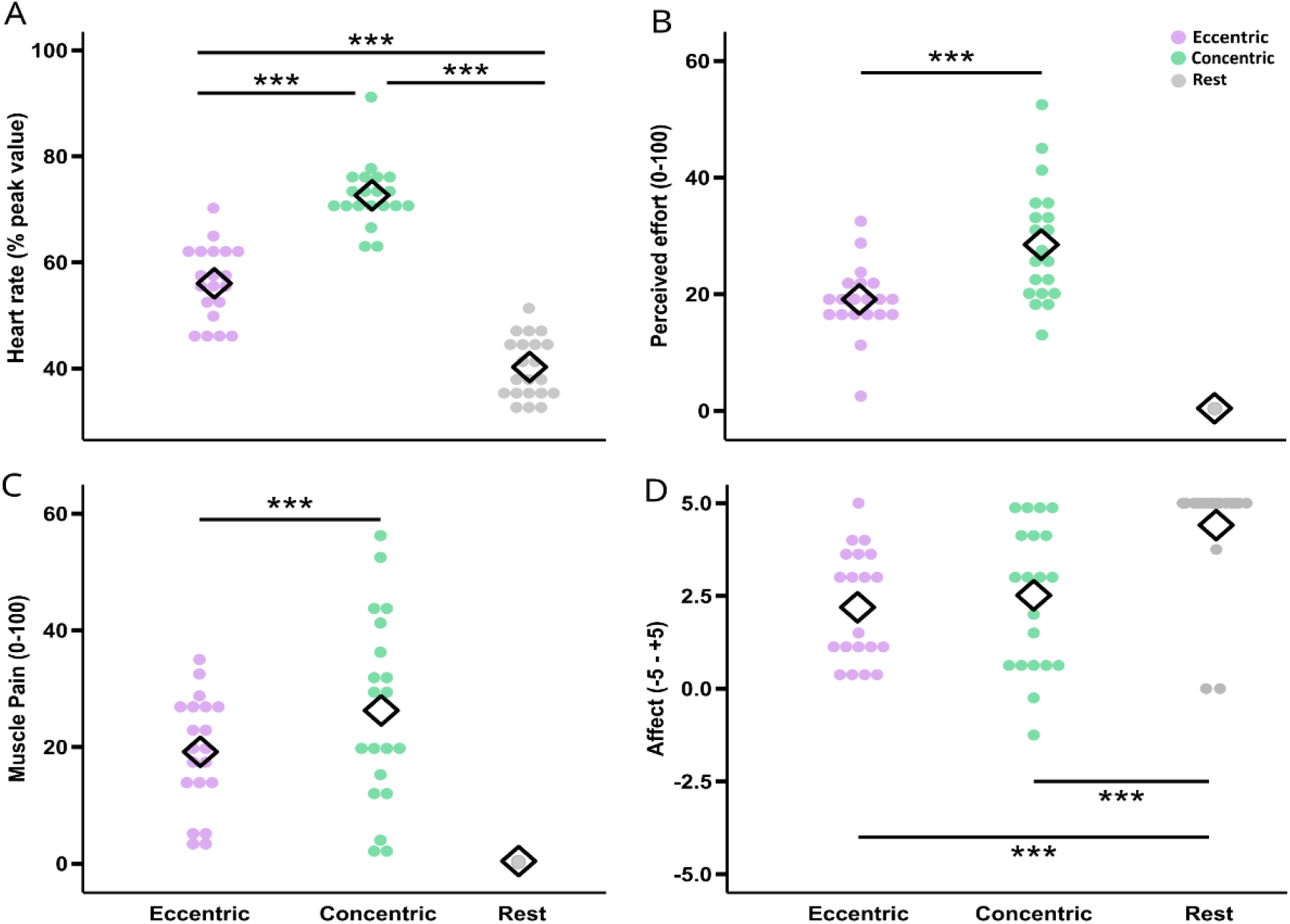
Parameters obtained during the intervention. All parameters are averaged over the 20-minute intervention. Panel A: Heart rate (% peak value). Panel B: perceived effort (CR100 Borg scale). Panel C: perceived muscle pain (CR100 Borg scale). Panel D: affect (Feeling scale from −5 to +5). Dots represent individual data points, while the black diamond represents the group mean. Purple dots refer to the eccentric cycling group, green dots to the concentric cycling group, and grey dots to the rest group. ***: *p*-value < .001.

*% HR*_*peak*_. There was a main effect of GROUP [*F*_*2,57*_ = 480.499, *p* = <.001, *η* ^*2*^_*p*_ *=* 0.811]. Post hoc analyses revealed that % HR_peak_ was higher for the concentric group compared to both eccentric (*p* < .001, *d* = −1.313) and rest (*p* < .001, *d* = −2.045) groups, and the eccentric group had a higher % HR_peak_ than the rest group (*p* < .001, *d* = −1.425). Additionally, there was no main effect of TIME [*F*_*3,171*_ = 0.753, *p* = .521, *η*^*2*^_*p*_ *=* 0.010], and no GROUP x TIME interaction [*F*_*6,171*_ = 1.284, *p* = .265, *η*^*2*^ _*p*_ *=* 0.033].

#### Perceived effort

There was a main effect of GROUP revealing that perceived effort was higher for the concentric group compared to the eccentric group [*F*_*1,38*_ = 34.379, *p* <.001, *η*^*2*^_*p*_ *=* 0.188]. Additionally, there was no main effect of TIME [*F*_*3,114*_ = 2.279, *p* = .081, *η*^*2*^_*p*_ *=* 0.044] and no GROUP x TIME interaction [*F*_*3,114*_ = 0.829, *p* =.479, *η*^*2*^_*p*_ *=* 0.016].

#### Muscle pain

There was a main effect of GROUP revealing that muscle pain was higher for the concentric group compared to the eccentric group [*F*_*1,38*_ = 9.523, *p* = .002, *η*^*2*^_*p*_ *=* 0.060]. Additionally, there was no main effect of TIME [*F*_*3,114*_ = 1.605, *p* = .191, *η*^*2*^_*p*_ *=* 0.031] and no GROUP x TIME interaction [*F*_*3,114*_ = 0.114, *p* = .951, *η*^*2*^_*p*_ *=* 0.002].

#### Affect

There was a main effect of GROUP [*F*_*2,57*_ = 37.030, *p* <.001, *η*^*2*^_*p*_ *=* 0.248]. Post hoc analysis revealed that affective responses for the rest group were higher compared to both concentric (*p* < .001, *d* = 1.084) and eccentric (*p* < .001, *d* = 1.369) groups, with no differences observed between both concentric and eccentric groups (*p* = .422 *d* = −0.127). Additionally, there was no main effect of TIME [*F*_*3,171*_ = 0.510, *p* = .675, *η*^*2*^_*p*_ *=* 0.007] and no GROUP x TIME interaction [*F*_*6,171*_ = .351, *p* = .910, *η*^*2*^_*p*_ *=* 0.009].

### Motor task performance

Figure 3 displays average RMSE data for each group (eccentric, concentric, rest) across all practice blocks, separately for acquisition and retention. Additionally, Figure S2 displays RMSE data with boxplots to show the individual variability of the data.

**Figure 3:**
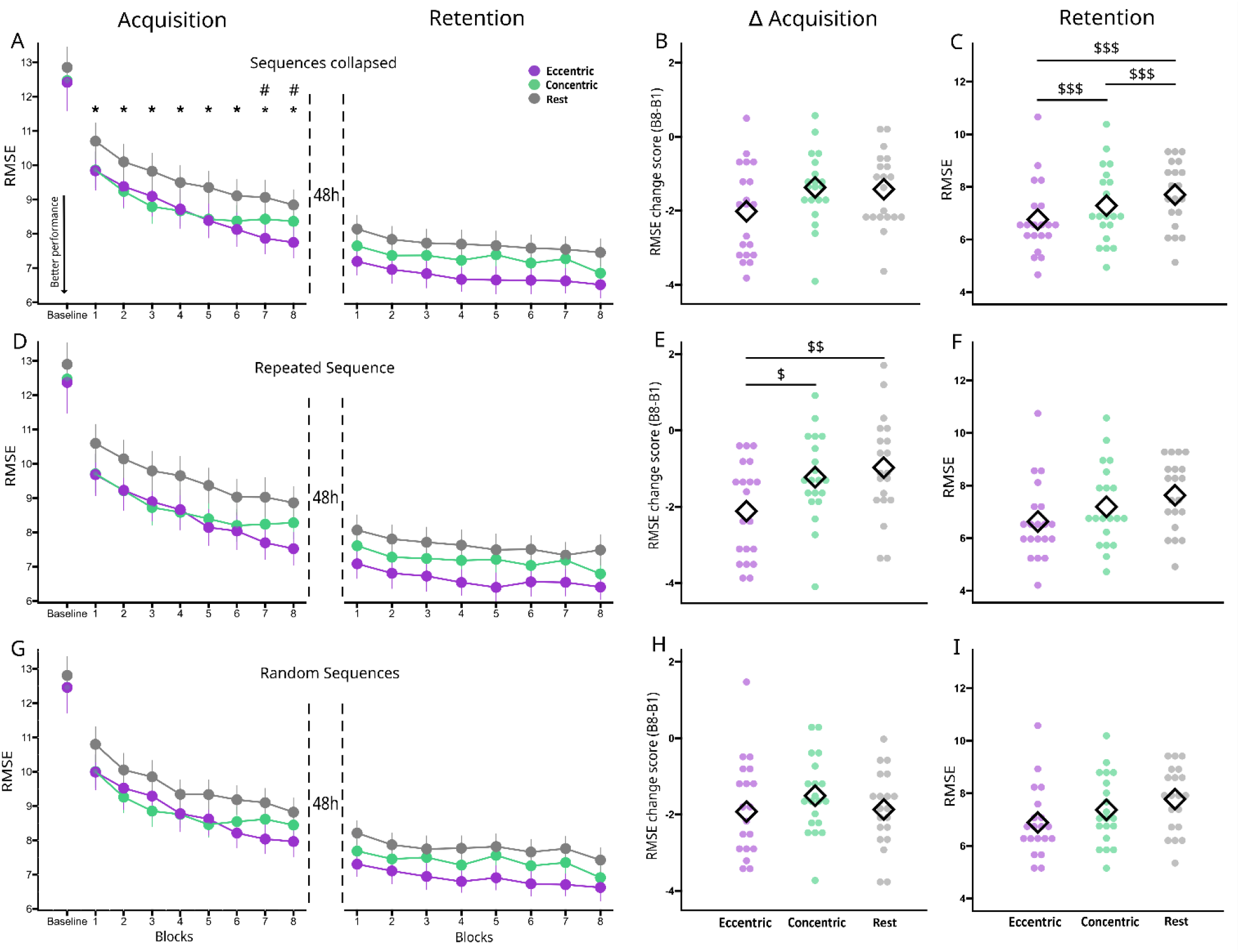
Motor learning data of the CTT. Panel A: Motor task performance when both repeated and random sequences are collapsed. Panel B: RMSE change score from Block 1 to Block 8 of acquisition when both repeated and random sequences are collapsed. Panel C: RMSE score of retention when both repeated and random sequences are collapsed. Panel D: Motor task performance for the repeated sequence. Panel E: RMSE change score from Block 1 to Block 8 of acquisition for the repeated sequence. Panel F: RMSE score of retention for the repeated sequence. Panel G: Motor task performance for the random sequences. Panel H: RMSE change score from Block 1 to Block 8 of acquisition for the random sequences. Panel I: RMSE score of retention for the random sequences. For panels A, D and G, data are averaged across the group and error bars represent standard error of the mean. **\***: significant difference at the block between both exercise groups (eccentric and concentric) and the rest group. ^**#**^: significant difference at the block between eccentric cycling group and both other groups (concentric and rest). For panels B, C, E, F, H and I, dots represent individual data points, while the black diamond represents the group mean. ^**$**^: *p*-value < .05; ^**$$**^: *p*-value < .01; and ^**$$$**^: *p*-value < .001. RMSE = root mean squared error; B8 = Block 8, B1 = Block 1; Δ = change score between Block 1 and Block 8. CTT = continuous tracking task. Purple dots refer to the eccentric cycling group, green dots to the concentric cycling group, and grey dots to the rest group. For all panels, a decrease in RMSE values refers to a better CTT performance.

#### Baseline measures

There was no main effect of GROUP [*F*_*2,57*_ = 0.724, *p* = .486, *η*^*2*^_*p*_ *=* 0.004], SEQUENCE [*F*_*1,57*_ = 0.188, *p* = .665, *η*^*2*^_*p*_ *=* 0.001], nor a GROUP x SEQUENCE interaction [*F*_*2,57*_ = 0.025, *p* = .975, *η*^*2*^_*p*_ *=*0.002].

#### Acquisition

There was a GROUP x BLOCK interaction (*F*_*14,399*_ = 2.242, *p* = .005, *η*^*2*^_*p*_ *=* 0.003]. Post hoc analyses showed differences between both exercise groups and rest (all *ps* < .018 all *ds* > 0.241) from blocks 1 to 8 with eccentric and concentric groups performing better than rest. However, at blocks 7 and 8, the eccentric group performed better than both the concentric group (Block 7: *p* = .004, *d* = −0.275; Block 8: *p* < .001, *d* = −0.304) and the rest group (Block 7: *p* < .001, *d* = −0.562; Block 8: *p* < .001, *d* = −0.543 ; [i.e., the end of acquisition; Figure 3A]). Additionally, we found a main effect of GROUP [*F*_*2,57*_ = 160.761, *p* < .001, *η*^*2*^_*p*_ *=* 0.033], and BLOCK [*F*_*7,399*_ = 18.959, *p* < .001, *η*^*2*^_*p*_ *=* 0.014], as well as a GROUP x SEQUENCE interaction [*F*_*2,57*_ = 4.059, *p* = .017, *η*^*2*^_*p*_ *=* 0.001]. However, we found no main effect of SEQUENCE [*F*_*1,57*_ = 0.429, *p* = .512, *η*^*2*^_*p*_ *=* 0.004].

#### Magnitude of CTT performance improvement during acquisition

Since our acquisition analysis showed differences between both exercise groups (eccentric and concentric) and the rest group at block 1, we followed up with further analysis examining whether eccentric or concentric AAE impacted the magnitude of RMSE reduction during acquisition. A change score between block 8 and block 1 (B8-B1) was performed to calculate the magnitude of RMSE change following AAE (eccentric, concentric) or rest (Figure 3B, 3E 3H). There was a GROUP*SEQUENCE interaction [*F*_*2,57*_ = 3.243, *p* = .039; *η*^*2*^_*p*_ *=* 0.008]. Post hoc analyses showed a greater RMSE decrease in the eccentric group compared to both concentric (*p* = .018, *d* = −0.280) and rest (*p* = .001, *d* = −0.317) groups for the repeated sequence, with no effects for the random sequences (all *ps* = .999 all *ds* < 0.176). Additionally, we found a main effect of GROUP [*F*_*2,57*_ = 5.808, *p* = .003, *η*^*2*^_*p*_ *=* 0.009], and a main effect of SEQUENCE [*F*_*1,57*_ = 6.042, *p* = .014; *η*^*2*^_*p*_ *=*.005].

#### Retention test

There was a main effect of GROUP [*F*_*2,57*_ = 234.497, *p* < 0.001, *η*^*2*^_*p*_ *=* 0.047]. Post hoc analyses revealed that the eccentric group performed better than both the concentric (*p* < .001, *d* = −0.297) and rest (*p* < .001, *d* = −0.537) groups, while the concentric group demonstrated better performance than the rest group (*p* < .001, *d* = −0.237; Figure 3C). Additionally, we found main effects of SEQUENCE [*F*_*1,57*_ = 6.702, *p* =.009, *η*^*2*^_*p*_ *=* 0.001] and BLOCK [*F*_*7,399*_ = 2.143, *p* = .036, *η*^*2*^_*p*_ *=* 0.002].

## Discussion

This study pioneers the investigation of acute eccentric cycling exercise’s impact on motor learning. We revealed that eccentric cycling significantly enhanced motor learning more than both concentric cycling and rest. Although concentric cycling amplified motor skill acquisition and learning relative to rest, eccentric cycling elicited a superior increase. Unexpectedly, eccentric cycling elicited increases in both sequence-specific and non-specific motor learning, as evidenced by performance gains observed during the 48-h retention test. Notably, while both exercise interventions (eccentric and concentric) yielded an immediate performance increase (reduced tracking error at block 1) compared to rest, eccentric cycling resulted in a continued reduction in tracking error. Further analysis revealed that eccentric cycling elicited a larger magnitude of sequence-specific motor skill acquisition immediately post-exercise, surpassing the effects of concentric cycling and rest. Our results suggest that priming motor learning with eccentric cycling offers an advantage over traditional concentric cycling exercise, which may have several implications in sport-related and clinical contexts. Here, we discuss the potential neural and behavioral mechanisms underlying the beneficial impact of eccentric cycling exercise on motor learning.

### Neural mechanisms of enhanced motor learning following eccentric exercise

Eccentric cycling enhanced both skill acquisition and motor learning, potentially by inducing neuroplastic adaptations that underlie the observed enhanced motor performance and learning. This is supported by findings from Latella et al. (2019), who found a decrease in intracortical inhibition (i.e., short-interval intracortical inhibition [SICI]) and an increase in intracortical facilitation immediately following both eccentric and concentric contractions ^30^. Notably, they also found a prolonged decrease in a unique circuit of M1 intracortical inhibition (i.e., long-interval intracortical inhibition) following eccentric compared to concentric contraction exercise ^30^. These findings suggest that eccentric exercise may induce long-lasting (up to 1 h post-exercise) modulations in cortical inhibitory circuitry, which may be a mechanisms underlying the motor performance and learning improvements observed following eccentric cycling. Our recent meta-analysis demonstrated that concentric cycling AAE consistently decreases inhibitory circuitry (e.g., SICI) within the non-exercised upper limb muscle representation in M1 ^53^. Other work has shown that other inhibitory circuitry (i.e., long-interval intracortical inhibition) is impacted following concentric cycling AAE (Mooney et al., 2016; Singh et al., 2014). Importantly, this reduced M1 inhibition has been shown to be driven by GABAergic activity ^54,55^, and has been associated with motor skill practice and learning ^34,56,57^. Taken together, it is possible that eccentric cycling exercise may have amplified and prolonged a reduction in M1 inhibitory circuitry compared to traditional concentric cycling, leading to the enhanced motor skill acquisition and motor learning we observed.

Another potential mechanism supporting the eccentric cycling exercise-enhanced motor learning observed may be via amplified corticospinal excitability as has been shown following concentric cycling ^58,59^. Previous research has shown that corticospinal excitability is modulated due to practicing motor tasks and that this excitability increase is associated with motor learning ^60-62^. Interestingly, neural mechanisms underlying eccentric contractions are complex, involving both supraspinal and spinal factors critical for motor control ^63^. Notably, at the cortical level, research suggests that eccentric movements are processed differently from concentric movements. Compared to concentric contractions, eccentric contractions generate greater cortical signals in sensorimotor-related cortical areas of the frontal and parietal lobe during movement planning and execution ^64^, and require longer preparation time and greater magnitude for execution ^65^. Relatedly, corticospinal excitability decreases before eccentric contractions but significantly increases following movement execution compared to concentric contractions ^66^. This shift in excitability during eccentric muscle contractions may suggest that eccentric cycling induces unique preparation and execution activity, leading to greater net increases in corticospinal excitability output. This may lead to a ‘priming’ of motor cortex excitability with repeated eccentric muscle contractions, resulting in greater capacity for neuroplastic change following eccentric cycling. This may have created a neural environment more susceptible for heighted corticospinal excitability change and neuroplasticity, which may have supported enhanced performance immediately post-exercise (skill acquisition), leading to enhanced motor learning of the CTT observed in our study. Yet, these points are speculative, and future research must be done to specifically investigate the proposed effects of eccentric cycling AAE on corticospinal excitability.

The increased activation of bilateral prefrontal cortex and right parietal lobe during eccentric cycling, as reported by Borot et al. (2024), provides a potential underlying neural mechanism to support the improved motor learning observed in our study ^31^. The prefrontal cortex plays a key role in cognitive, explicit and executive processes ^32^, which are important functions during the early cognitive stage of motor learning ^67^. The right parietal lobe is involved in sensorimotor integration and movement planning, critical for motor skill acquisition. The greater engagement of the bilateral prefrontal cortex and right parietal lobe during eccentric cycling suggests that this exercise modality may facilitate motor learning by enhancing the neural processes underlying motor planning, sensorimotor integration and movement refinement. These findings support the idea that eccentric cycling engages cognitive functions that may create a favorable neural environment for motor learning.

### Behavioral mechanisms of enhanced motor learning following eccentric exercise

The enhanced motor learning observed after eccentric cycling may stem from two interconnected behavioral mechanisms: 1) task familiarization and learning, and 2) heightened cognitive demand. First, eccentric cycling may require a learning process, as participants adapt to resisting backward pedal motion while maintaining a target cadence ^68-70^. Studies vary on the required familiarization time, ranging from a single 15-minute session to multiple 20-minute sessions ^31,68,71,72^. Previous research suggests that multiple 20-minute sessions of eccentric cycling practice are necessary to reduce cadence variability ^68^, with performance continuing to improve until the 6th or 8th session, indicating a task-learning process associated with eccentric cycling ^68^. However, other studies have used a single ∼15-minute familiarization session of eccentric cycling, suggesting that this may be sufficient for performance stabilization ^31,71,72^. Therefore, the eccentric familiarization in the current study may have been adequate to attain stabilization of cycling performance. Taken together, it is possible that eccentric cycling provided a priming effect on our target motor task (i.e., the CTT) via familiarization and learning the task of eccentric cycling itself, though further research is needed to determine if this constitutes motor skill learning and its impact on subsequent tasks.

Second, eccentric cycling may be more cognitively demanding than concentric cycling, which could enhance motor learning of a subsequent motor task (e.g., the CTT). Evidence shows eccentric cycling is more mentally demanding than concentric cycling, and engages increased frontal and parietal brain activity ^31^. This aligns with findings that cognitively challenging physical exercise, such as immersive virtual reality training or team sports, improves cognitive-motor performance and executive function ^73-75^. Combining physical and cognitive tasks can also yield greater benefits than either alone ^74^. Therefore, the higher cognitive demand of eccentric cycling may have contributed to the observed motor learning improvements, especially during the early, cognitively demanding stages of learning ^76-80^. However, further research is necessary to fully understand the cognitive demands of eccentric cycling and its influence on learning a subsequent target motor task.

### Practical implications

Incorporating eccentric cycling into training and rehabilitation programs may be useful to enhance motor learning, potentially making it a valuable tool for both athletes and individuals undergoing rehabilitation. For athletes, eccentric cycling could serve as a complementary training method to improve coordination and enhance motor performance, particularly in disciplines requiring fine motor control and neuromuscular precision. In clinical populations, eccentric cycling may be useful as an adjunct intervention to aid motor function recovery, since frontoparietal network activation is linked to motor recovery as demonstrated in ischemic stroke patients (Olafson et al., 2022). Additionally, eccentric cycling is feasible for patients with coronary heart disease ^27^ and its low metabolic cost makes it particularly suitable for those with limited cardiovascular capacity. Eccentric cycling may offer a unique advantage by promoting motor learning during rehabilitation sessions without a high cardiovascular demand, as many patients (e.g., cardiovascular, ischemic stroke) are often deconditioned and may have limitations to perform higher intensities of exercise.

Our findings also show that perceived effort and muscle pain were lower during eccentric cycling compared to concentric cycling, further supporting its suitability for clinical populations. These characteristics, combined with its superior effects on motor learning, highlight its potential in rehabilitation settings. By offering an exercise modality which minimizes effort and metabolic cost, while potentially enhancing motor learning and control, eccentric cycling may be useful as an adjunct in rehabilitation interventions for a wide range of populations.

## Limitations

Our study has some limitations. First, while our study matched the power output between conditions, physiological demands (e.g., VO_2_ or heart rate) were not. Future research could address this limitation by matching physiological demands across conditions to better understand the specificity of eccentric cycling in priming motor learning. Second, the exercise intensity used in the current study was based on measured HR_peak_, acknowledging that this method may be less precise and more variable than alternatives like oxygen uptake (VO2) or HR reserve. Our results should be conceptually reproduced using different exercise intensity prescription methods to confirm their consistency. Third, muscle pain was only assessed during the intervention and not beyond the exercise intervention, despite eccentric cycling often being associated with delayed onset muscle soreness ^70,81^. Future studies could include assessments up to 96 hours post-exercise to evaluate whether the benefits of eccentric cycling on motor learning are worth the muscle pain elicited after eccentric cycling. Finally, the results of this study may not be generalizable to other populations since the sample used in this study does not represent diverse age groups, fitness levels, or clinical populations, limiting the broader applicability of the findings.

## Conclusion

Our study reveals that eccentric cycling leads to greater improvement in motor learning compared to traditional concentric cycling and rest. The enhanced motor learning observed in the eccentric cycling group may be attributed to the unique neurophysiological and/or behavioral adaptations elicited by repeated eccentric contractions. Our findings may have implications as a promising avenue for enhancing motor learning in sports and athletic training contexts as well as enhancing the efficacy of rehabilitation programs. This research paves the way for further exploration into the benefits of eccentric cycling exercise, potentially leading to more effective and innovative strategies for improving motor performance and recovery of function in clinical populations.

## Author contributions

LY contributed to developing the study design, data collection and analysis, and wrote the first draft of the manuscript. JLN and BP conceived the study and contributed to developing the study design, data interpretation, and participated in writing and editing the manuscript. All authors read and approved the final version and agreed on the authorship order.

## Data availability

The datasets generated and analyzed during the current study are available from the corresponding author upon reasonable request, and approval of the ethics committee. Individual data are presented in the figures.

## Supplementary material

**Figure S1:**
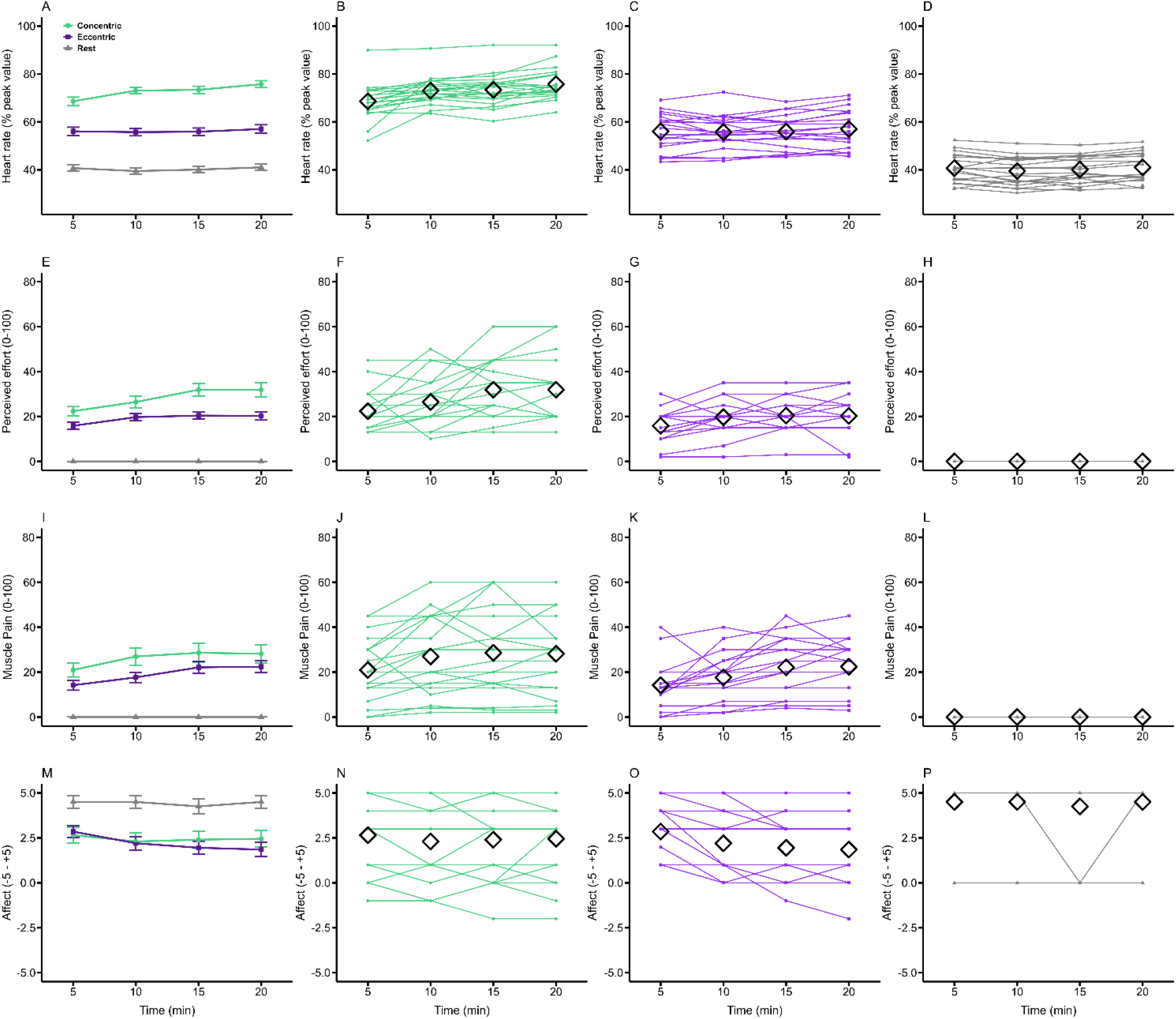
Parameters obtained across the intervention. Green represents concentric group, purple represents eccentric group, and grey represents rest group. Time points of 5, 10, 15, and 20 minutes indicate when parameters were recorded during the intervention. The black diamond indicates the average at each time point. *Heart rate (% peak value):* Panel A shows the averages for each group, while Panels B, C, and D display individual data points for the concentric, eccentric, and rest groups respectively. *Perceived effort:* Panel E shows the averages for each group, while Panels F, G, and H display individual data points for the concentric, eccentric, and rest groups, respectively. *Muscle pain:* Panel I shows the averages for each group, while Panels J, K, and L display individual data points for the concentric, eccentric, and rest group, respectively. *Affect:* Panel M shows the averages for each group, while Panels N, O, and P display individual data points for the concentric, eccentric, and rest group, respectively.

**Figure S2:**
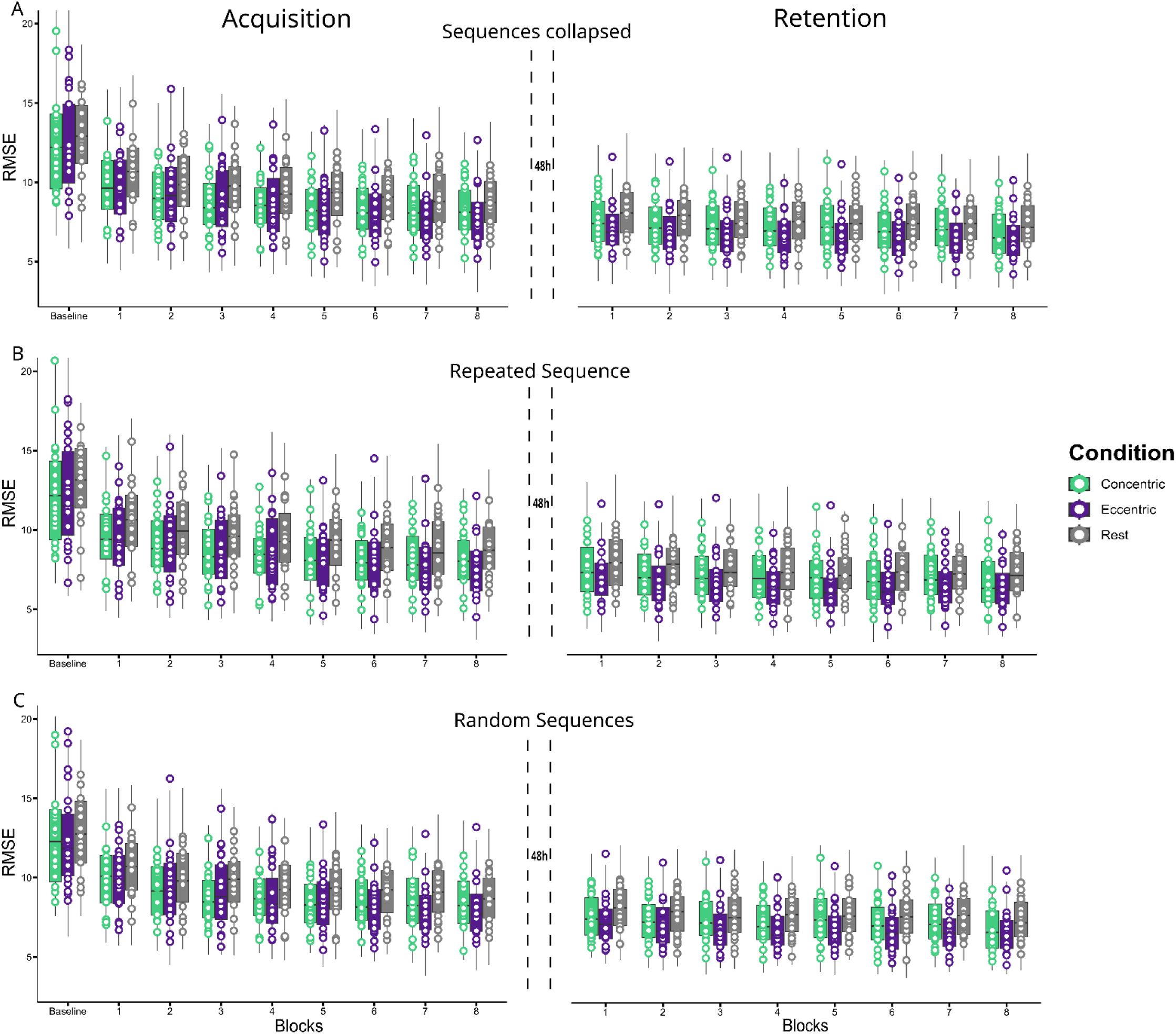
Motor learning data of the CTT visualising individual data. Panel A: Motor task performance of the individual data when both repeated and random sequences of the CTT are collapsed for both acquisition and retention. Panel B: Motor task performance of the individual data for the repeated sequence of the CTT for both acquisition and retention. Panel C: Motor task performance of the individual data for the random sequences of the CTT for both acquisition and retention. Purple refers to eccentric cycling group, green refers to concentric cycling group, and grey refers to rest group. RMSE = root mean squared error; CTT = continuous tracking task.

